# First concrete documentation for presence of *Aedes (Stegomyia) albopictus* in Bolivia: Dispelling previous anecdotes

**DOI:** 10.1101/2024.03.14.585027

**Authors:** Frédéric Lardeux, Philippe Boussès, Rosenka Tejerina-Lardeux, Audric Berger, Christian Barnabé, Lineth Garcia

## Abstract

**Background:** The presence of *Aedes albopictus* in Bolivia has been a subject of controversy, lacking concrete documentation until now.

**Objectives:** This study aimed to furnish evidence of *Ae. albopictus* presence in Bolivia.

**Methods:** Larval breeding sites were sampled in two northern Bolivian localities, Rosario del Yata and San Agustin, both situated in the Beni department within the Vaca Diez province and Guayaramerin Municipio, approximately 10 km apart. Mosquito larvae collected underwent rearing to L4 and adult stages for morphological identification, with some specimens sequenced for confirmation.

**Findings:** *Ae. albopictus* was identified in multiple breeding sites in both localities, confirming its establishment in the area. This marks the first concrete documentation of the species in Bolivia. The collections (larvae and adults) have been deposited in the Medical Entomology Laboratory of the Universidad Mayor de San Simón in Cochabamba, Bolivia, and the Laboratory of Entomology of the Instituto Nacional de Laboratorios de Salud of the Ministry of Health in La Paz, Bolivia.

**Main conclusion:** Acknowledging its role as a vector for arboviruses like dengue and Chikungunya, *Ae. albopictus* should be incorporated into the Bolivian National Program of Vector Control for monitoring.

## INTRODUCTION

*Aedes (Stegomyia) albopictus* (Skuse, 1895) (*Stegomyia albopicta* sensu Reinert et al. (^1^)) (Diptera: Culicidae) is considered by the Invasive Species Specialist Group (ISSG) as one of the top 100 invasive species (^2^) and certainly the most invasive mosquito in the world. Commonly known as the ‘tiger mosquito’ and originally from tropical forests of South-East Asia, it has experienced a swift global expansion over the past four decades, propelled by human activities (^3^). The initial sighting outside its original distribution range occurred in 1979 in Albania (^4^), and presently it has established a global presence on all continents, excluding Antarctica (^5^).

In Latin America, *Ae. albopictus*, following its initial discovery in Brazil in 1986 (^6^), was subsequently documented in Mexico in 1988. In the period from 1993 to 1998, its presence extended to the Dominican Republic, Cuba, Guatemala, the Cayman Islands, Colombia, and Argentina. Throughout the 2000s, the mosquito further disseminated to Canada, Bermuda, Trinidad and Tobago, Panama, Nicaragua, Costa Rica, Uruguay, Venezuela, Belize, and Haiti. More recent reports indicate its identification in Ecuador in 2017 and Jamaica in 2018 (^7^).

Bolivia, located in central South America, is a landlocked country, bordered by Brazil to the north and east, Paraguay and Argentina to the south, Chile to the southwest, and Peru to the northwest. While Brazil and Argentina have documented the presence of *Ae. albopictus* within their territories, there is currently no indication of the mosquito being reported in Chile, and Peru (^7^). In Paraguay, the species has likely been found in various departments, including one bordering Bolivia (Boquerón)(^8^), albeit never officially published in a scientific article (^7^). In Argentina, *Ae. albopictus* has been identified exclusively in the two provinces of Misiones and Corrientes, situated in the northwest of the country, bordering Paraguay. Notably, there is a considerable distance from the Bolivian border, approximately 1000 km, and no records of its presence have been reported in the intervening regions (^9^). In Brazil, the states bordering Bolivia, specifically Rondônia, Mato Grosso, and Mato Grosso do Sul, reported the presence of *Ae. albopictus* between 1991 and 2002 (^10^). More recent observations confirming the presence of the mosquito have come from Acre state in 2022, specifically in the city of Rio Branco, located approximately 70 km in a straight line from the Bolivian border (^11^).

The presence of *Ae. albopictus* in Bolivia has been a subject of controversy. Initial scientific reports were solely reliant on personal communications, particularly from former PAHO officials who held responsibilities in Bolivia before the 2000s ^(12; 13)^. These reports indicated the species’ presence in the country since 1997 or potentially slightly earlier (^14^), lacking further details. However, this presence was not confirmed in the subsequent PAHO report (^15^). Nevertheless, these personal communications stem from an anecdote recounted to one of the authors of this article (FL) which mentioned that in the 1990s, *Ae. albopictus* was reportedly collected in Cotoca in the Santa Cruz Department without specifying whether “Cotoca” referred to the Botanical Garden of the city of Santa Cruz or a locality within the Department. The sample was reportedly sent to a laboratory in Argentina for species identification confirmation, yet neither the specific laboratory nor the contact scientist was disclosed. Although the identification was purportedly set to be confirmed, the sample was regrettably misplaced, and all subsequent efforts to locate it have proven futile, casting doubts on the actual existence of the collected specimens. Recently, in March 2023, a team of entomologists based in Santa Cruz de la Sierra claimed to have discovered the species in four locations within the Municipio de San Ignacio de Velasco (Santa Cruz Department, Province of José Miguel de Velasco) (Lat. -16.38, Long. -60.9) as reported in a local press article (^16^) and via social networks. (^17^). Unfortunately, as far as we are aware, none of the leading entomological laboratories in Bolivia, including those in La Paz (INLASA) and Cochabamba (LEMUMSS), as well as the Ministry of Health, have received samples for confirmation, despite their persistent requests. Once again, legitimate doubts persist regarding the existence of *Ae. albopictus* specimens and/or their accurate identification.

The three databases, (1) the GBIF database, encompassing 90 413 worldwide references (^18^), (2) the WRBU-VectorMap database and (3) the “Global compendium of *Aedes aegypti* and *Ae. albopictus* occurrence” ^(5; 19)^, all lack records of *Ae. albopictus* occurrences in Bolivia. Bibliographic searches using standard databases (Web of Science, PubMed, bioRxiv) do not reveal any documented occurrences of the species in Bolivia. In fact, there is no document or voucher conclusively demonstrating the presence of *Ae. albopictus* in the country (^7^).

The present article details the first documented discovery of *Ae. albopictus* in two localities in northern Bolivia, Rosario del Yata and San Agustin, within the Beni Department (Provincia Vaca Diez, Municipio Guayaramerin).

## MATERIAL AND METHODS

### Collection of mosquitoes

*Ae. albopictus* specimens were collected during an entomological survey conducted from November 12 to December 4, 2023, in the Amazonian region of Bolivia as part of the VECTOBOL initiative (https://vectobol.ird.fr/), jointly organized by Institut de recherche pour le développement (IRD, France) and Instituto de Investigaciones Biomédicas e Investigación Social (IIBISMED, Bolivia).

The mission was carried out by three of the article’s authors (FL, PB, and RTL) in search of wild Culicidae in the Amazonian forest environment. However, passing through the Rosario del Yata (lat. -10.99, long. -65.58) on November 23, 2023, a locality situated about 30 km west of Guayaramerin, along the Guayaramerin-Riberalta road, the team sampled a 200-liter plastic container collecting rainwater in the peridomicile of a house (sample ID: FLX-2023-021 of Table 1). The collection mainly consisted of numerous small larvae resembling the genus “*Aedes*” and some *Toxorhynchites*. To precisely identify the species, larvae were reared to the L4 and adult stages. Subsequently, the first *Ae. albopictus* adults were morphologically identified (by RTL) and confirmed (by PB and FL). After this discovery, it was decided to conduct intensive sampling of the peridomiciles in Rosario del Yata. This sampling occurred on November 28, concluding the sampling carried out on November 26. Moreover, to verify the species’ establishment in this region, the adjacent locality of San Agustin (lat. -10.95, long. -65.49), approximately ten kilometers to the east, was sampled on November 27 (Fig 1 and Fig. 2).

**Table 1.** Samples and mosquito species collected in Rosario del Yata and San Agustin.

| Locality | Environment | Sample ID in laboratory | Type of collection* | Collecting site | Date of collection | Collected species |
| --- | --- | --- | --- | --- | --- | --- |
| Rosario del Yata** | Peridomestic | FLX-2023-021 *** | L | 200-l plastic drum | 23/11/2023 | *Ae. albopictus*<br>*Ae. aegypti*<br>*Tx. haemorrhoidalis* |
| Rosario del Yata | Peridomestic | FLX-2023-042 | L | Used tire | 28/11/2023 | *Ae. aegypti* |
| Rosario del Yata | Peridomestic | FLX-2023-043 | L | Cut plastic bottle with citric cutting in water | 28/11/2023 | *Ae. albopictus* |
| Rosario del Yata | Peridomestic | FLX-2023-044A | L | Cut plastic bottle with rain water | 28/11/2023 | *Ae. albopictus* |
| Rosario del Yata | Peridomestic | FLX-2023-045 | L | Tire animal waterer | 28/11/2023 | *Cx. coronator* |
| San Agustin | Peridomestic | FLX-2023-032 | L | 100-l concrete animal waterer | 27/11/2023 | *Cx. coronator*<br>*Cx. quinquefasciatus*<br>*Cx. corniger* |
| San Agustin | Peridomestic | FLX-2023-032B | L | Cut plastic bottle | 27/11/2023 | *Cx. quinquefasciatus* |
| San Agustin | Peridomestic | FLX-2023-032C | L | Aluminum pot | 27/11/2023 | *Ae. aegypti* |
| San Agustin | Peridomestic | FLX-2023-033A | L | 200-l plastic drum | 27/11/2023 | *Ae. aegypti* |
| San Agustin | Peridomestic | FLX-2023-033B | L | Plastic bucket | 27/11/2023 | *Ae. albopictus*<br>*Ae. aegypti*<br>*Tx. haemorrhoidalis* |
| San Agustin | Peridomestic | FLX-2023-033C | L | 10-l plastic canister | 27/11/2023 | *Ae. albopictus*<br>*Ae. aegypti*<br>*Cx. declarator* |
| San Agustin | Peridomestic | FLX-2023-034 | L | Used tire | 27/11/2023 | *Cx. declarator* |
| San Agustin | Peridomestic | FLX-2023-035 | L | 1000-l plastic drum | 27/11/2023 | *Ae. fluviatilis*<br>*Cx. quinquefasciatus* |
| San Agustin | Peridomestic | FLX-2023-036 | L | 200-l plastic drum | 27/11/2023 | *Ae. fluviatilis*<br>*Cx. quinquefasciatus* |
| San Agustin | Peridomestic | FLX-2023-037 | L | 200-l plastic drum | 27/11/2023 | *Ae. fluviatilis*<br>*Cx. quinquefasciatus* |
| San Agustin | Peridomestic | FLX-2023-038A | L | Plastic bathtub | 27/11/2023 | *Cx. quinquefasciatus* |
| San Agustin | Peridomestic | FLX-2023-038B | L | Tire animal waterer | 27/11/2023 | *Ae. fluviatilis* |
| San Agustin | Peridomestic | FLX-2023-039 | L | Tire animal waterer | 27/11/2023 | *Ae. fluviatilis*<br>*Ps. albigena* |
| San Agustin | Peridomestic | FLX-2023-040 | L | 30-l plastic bucket | 27/11/2023 | ***Ae. albopictus***<br>*Ae. aegypti* |
| Rosario del Yata | Peridomestic | FLX-2023-044B | A | Human bait catches | 28/11/2023 | ***Ae. albopictus***<br>*Ma. titillans* |
| Rosario del Yata | Riverbank | FLX-2023-019A | L | water in the bottom of a small wooden boat | 23/11/2023 | *Ae. fluviatilis*<br>*Cx. coronator*<br>*Culex. sp.* |
| Rosario del Yata | Riverbank | FLX-2023-019B | L | water in the bottom of a small wooden boat | 23/11/2023 | *Cx. coronator*<br>*Cx. glyptosalphinx* |
| Rosario del Yata | Riverbank | FLX-2023-019C | L | water in the bottom of a small wooden boat | 23/11/2023 | *Cx. coronator* |
| Rosario del Yata | Riverbank | FLX-2023-019D | L | In the river (riverside) | 23/11/2023 | *Cx. pilosus*<br>*Anopheles sp.* |
| Rosario del Yata | Riverbank | FLX-2023-020 | L | Hole in sandy bank with water infiltration | 23/11/2023 | *Ps. albigena* |
| Rosario del Yata | Close to forest | FLX-2023-022A | L | 1000-l plastic drum | 23/11/2023 | *Ae. fluviatilis*<br>*Cx. declarator* |
| Rosario del Yata | Close to forest | FLX-2023-022B | L | 1000-l plastic drum | 23/11/2023 | *Cx. declarator*<br>*Tx. haemorrhoidalis* |
| Rosario del Yata | Close to forest | FLX-2023-022C | L | 200-l plastic drum | 23/11/2023 | *Ae. fluviatilis*<br>*Cx. coronator* |
| Rosario del Yata | Close to forest | FLX-2023-022D | A | Human bait catches | 23/11/2023 | *Ma. titillans* |

\*Type of collection: L = larvae, A= adults (on human bait)
\*\*Locality where *Ae. albopictus* was first discovered;
\*\*\*First record of *Ae. albopictus*

**Figure 1.**
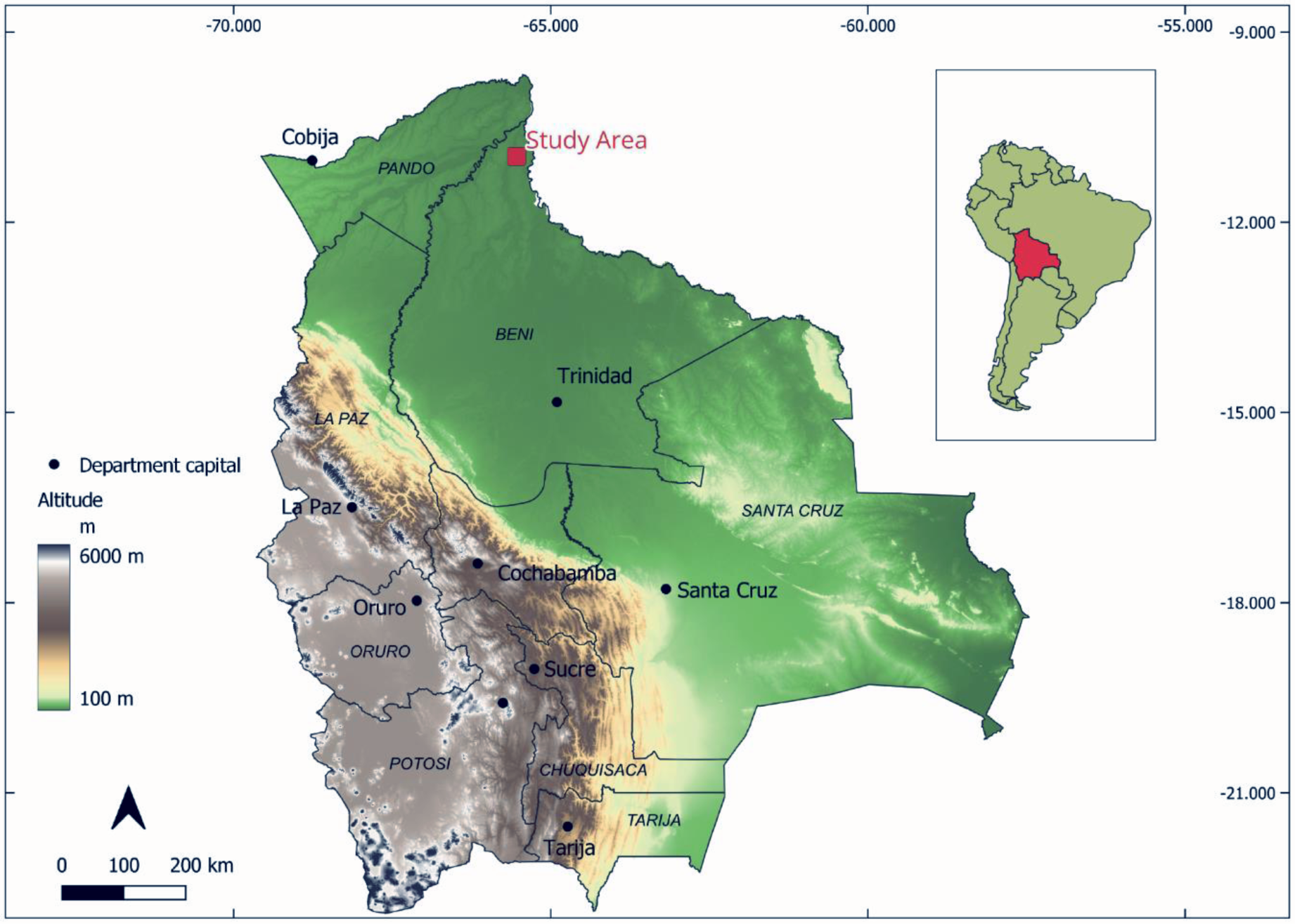
Location of the study area within Bolivia, situated in the Amazonian region near the border with Brazil.

**Figure 2.**
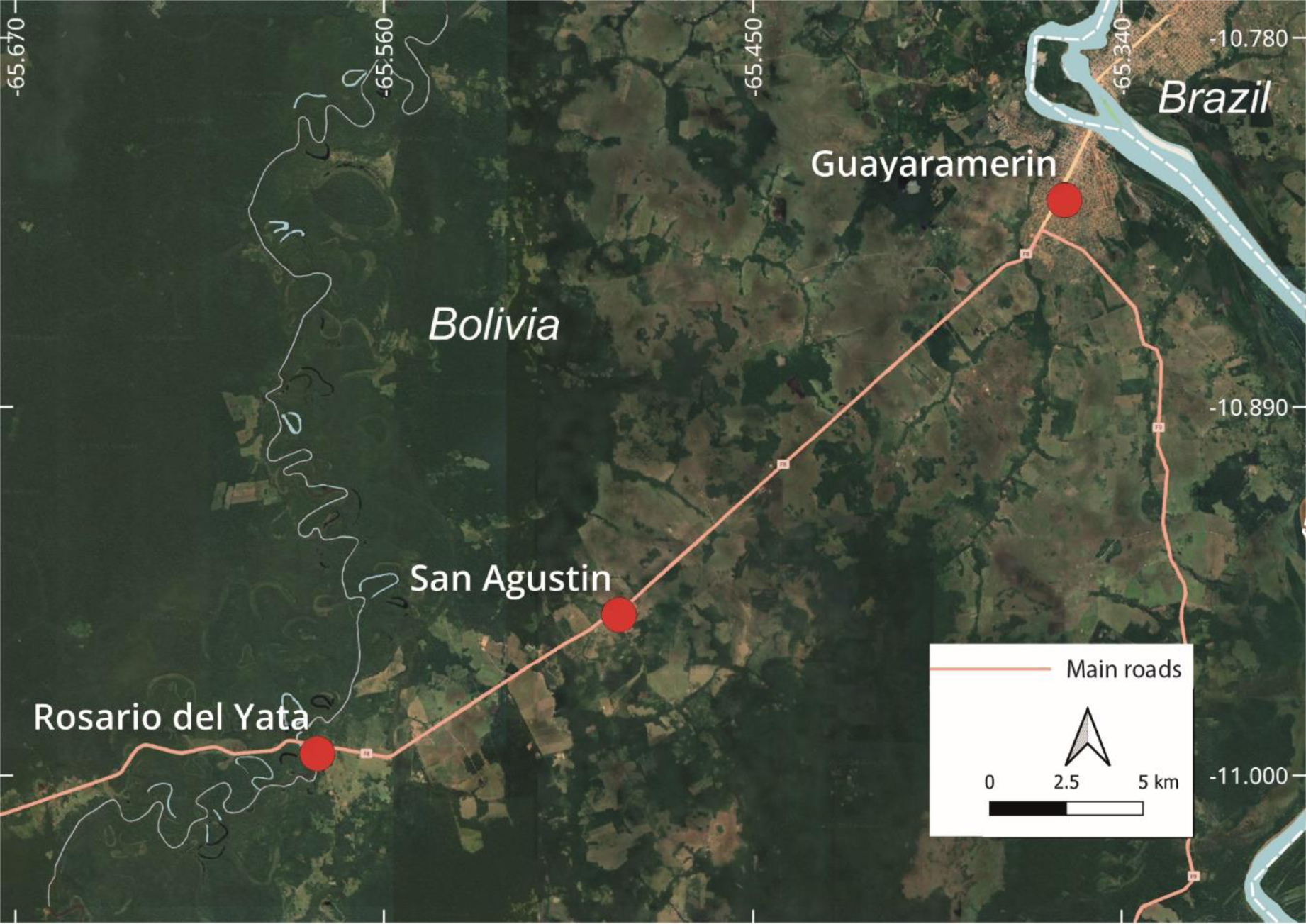
Study area: Localization of the two localities, Rosario del Yata and San Agustin, situated along the main road from Guayaramerin (bordering Brazil) to Riberalta, in the Bolivian Amazonian region.

Rosario del Yata is home to approximately 700 inhabitants residing in 180 houses, while San Agustin accommodates around 70 inhabitants in 15 houses. Both localities are located along the main road between the two major cities, Guayaramerin and Riberalta, and are in close proximity to the Amazonian forest that borders the road. Houses are modest, lacking fences for separation. The backyards are kept relatively neat and include water wells. Due to the tropical climate, people engage in domestic activities outside their houses, often leaving small containers like buckets, pans, and pots outdoors, which serve as potential breeding sites for mosquito larvae. Domestic animals primarily include dogs, ducks, and chickens, provided water through waterers generally made from halved tires cut longitudinally. Rainwater is also collected from roofs in 200-liter drums or other small containers. Due to proximity to the Amazonian forest, houses are amidst a tree-lined environment with trees that offer shade (Fig. 3)

**Figure 3.**
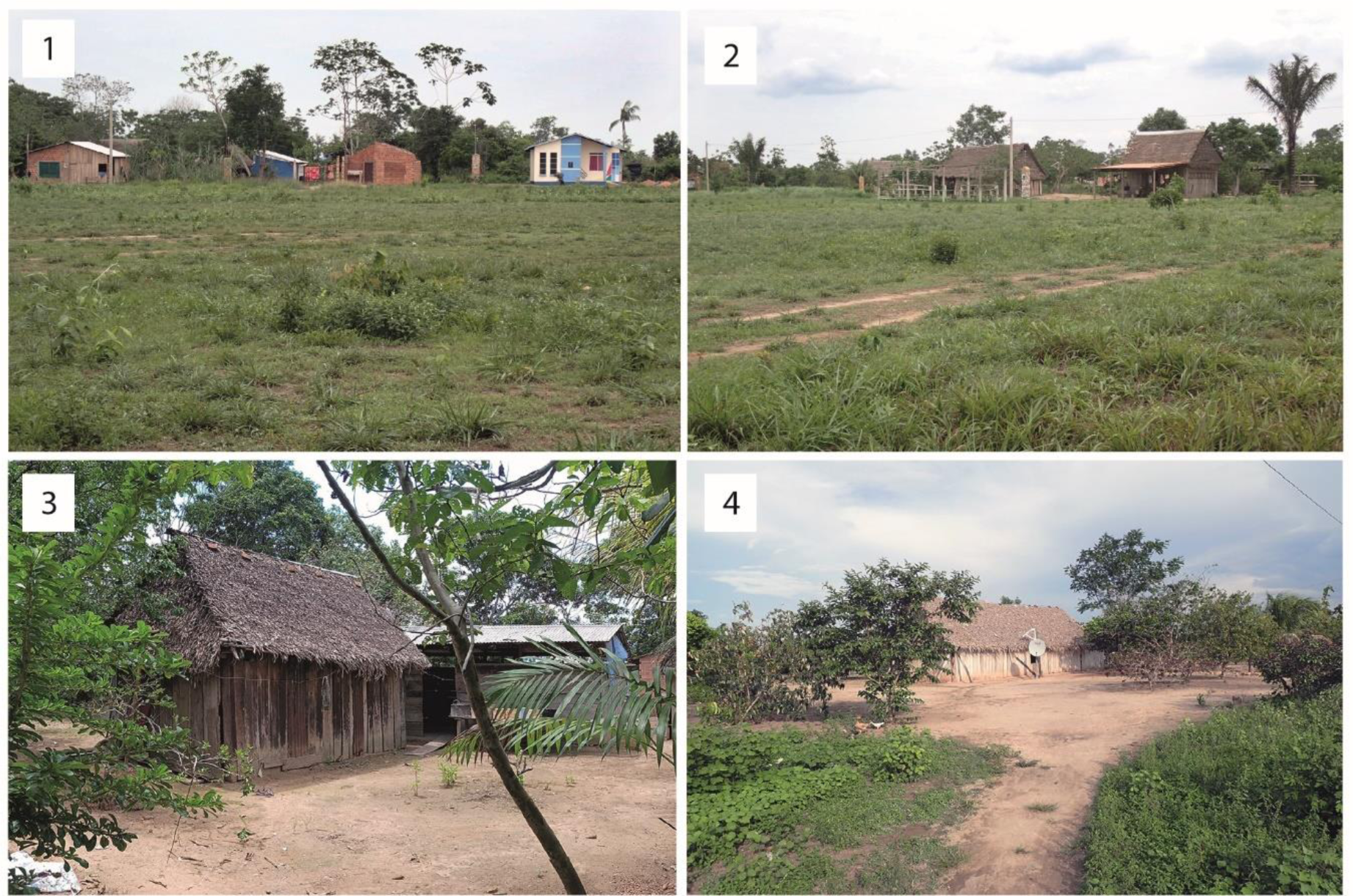
General view of human dwellings in Rosario del Yata (1, 2) and San Agustin (3, 4) depicting the dispersed habitat amidst a tree-lined environment.

In both localities, larval breeding sites in the peridomicile primarily comprised artificial recipients such as 200-liter drums to collect rain water from roofs, old tires used for domestic animal waterer, waste products such as plastic bottles, buckets etc. (Table 1) In Rosario del Yata, three types of environment were sampled: (1) the peridomestic environment where sampling was conducted in five human dwellings resulting in the collection of five larval samples (Table 1); (2) the “river bank environment “where samples consisted in the previous samples taken four days earlier near the Yata River, which borders Rosario del Yata. There, the collection points included the riverside, the bottom of small wooden boats, and a hole formed by water infiltration along the steep river bank (five samples, ID: FLX-2023-019A to C and FLX-2023-020 in Table 1); and (3) a “close to forest environment” which consisted in a small hangar used for walnut oil extraction, located approximately 2.5 km straight-line distance from Rosario del Yata in the Amazonian forest (four samples, ID: FLX-2023-022A to D in Table 1). All of these additional samples offered valuable insights into the Culicidae fauna present in the area.

In San Agustin sampling was conducted in nine human dwellings, resulting in the collection of 14 larval samples in a peridomestic environment (Table 1).

The locations of the samples are depicted in Fig. 4 for Rosario del Yata and Fig. 5 for San Agustin. Specific characteristics of the samples are detailed in Table 1. In all sampled breeding sites, larvae were reared to the L4 instar and adult stages before preservation, utilizing 70% alcohol for larvae and pinning for adults.

**Figure 4.**
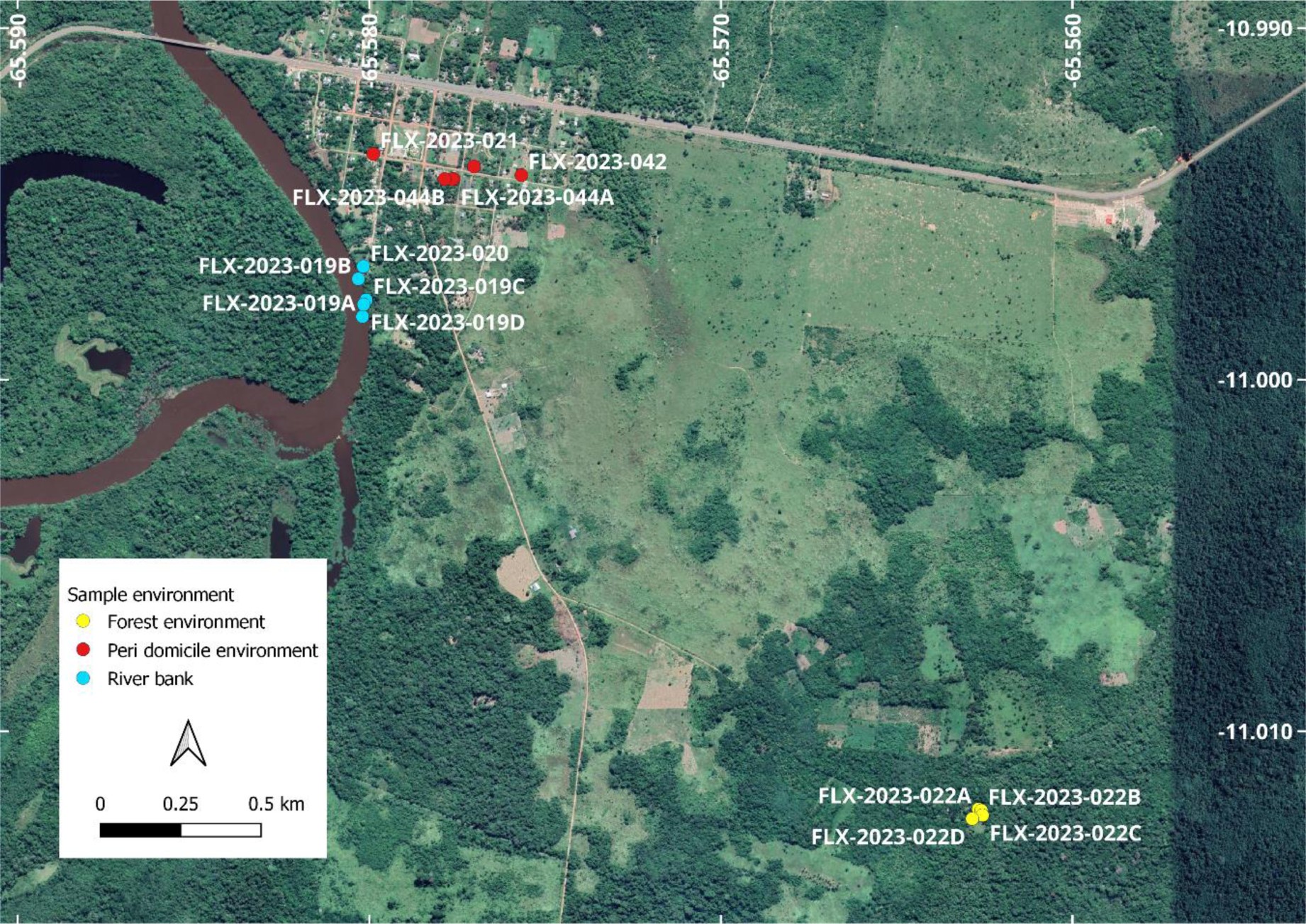
Localization of the sampling points in Rosario del Yata.

**Figure 5.**
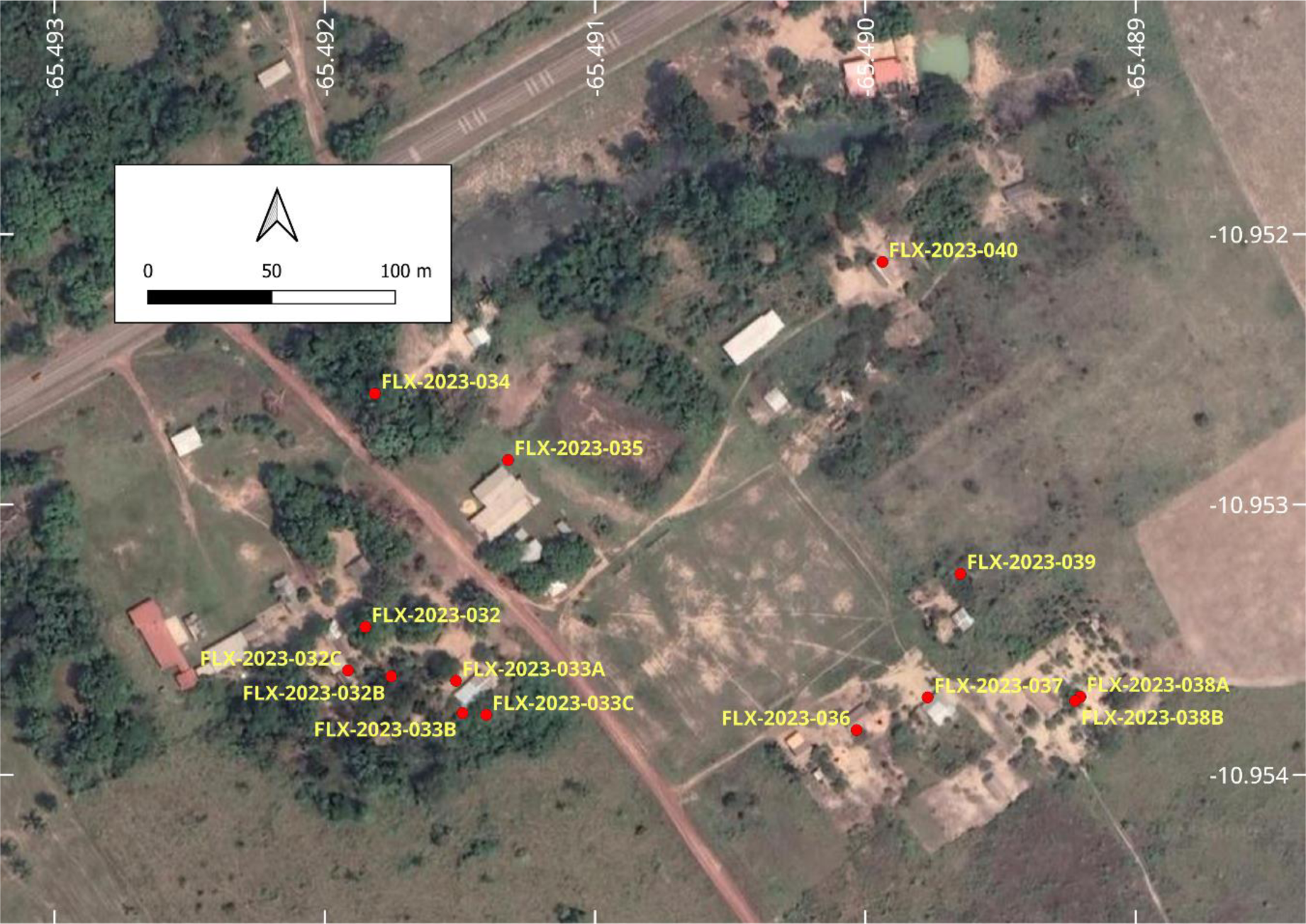
Localization of the sampling points in San Agustin.

### Morphological identification

*Aedes albopictus* was morphologically identified and differentiated from the unique other *Stegomyia* species of Bolivia, *Ae. aegypti*, using standard identification keys ^(20; 21; 22)^ and princeps or other descriptions ^(23; 24; 25; 26)^, from which distinctive diagnostic characters were extracted for adults (Fig. 6) and larvae (Fig. 7). Others species were identified using general keys ^(21; 27)^ and princeps or other descriptions of the species.

**Figure 6.**
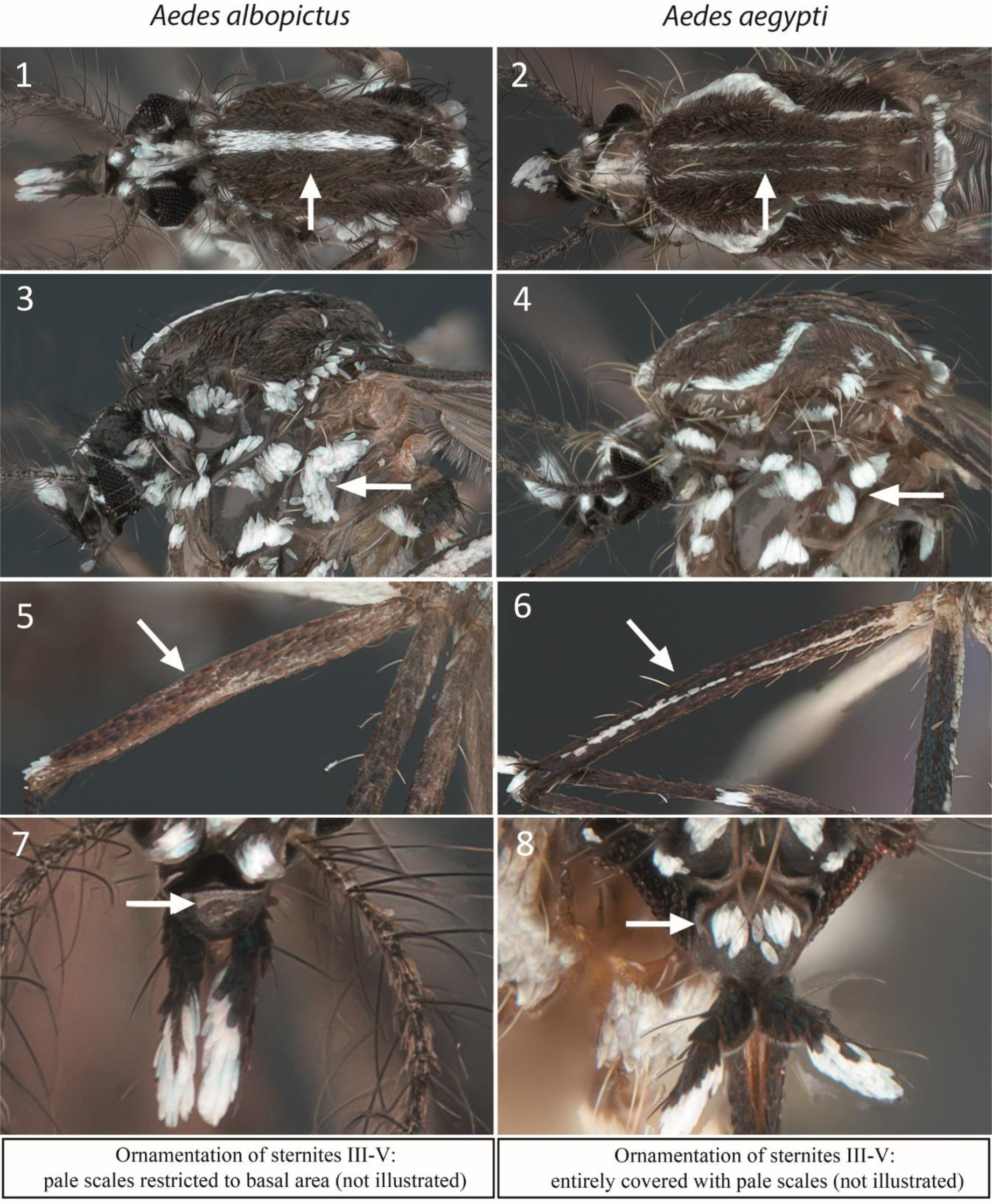
Morphological characteristic used to distinguish between *Ae. albopictus* and *Ae. aegypti* in adults. Specimens are from sample FLX-2023-021 of Rosario del Yata. Ornamentation of the scutum: narrow white median longitudinal stripe against a black background in *Ae.* albopictus (1), lyre-shaped silvery white stripes and narrow submedian longitudinal white stripes against a black background in *Ae. aegypti* (2). Ornamentation of the mesepimeron: two white scales spots connected in a unique V shape spot in *Ae albopictus* (3), two white scales spots well separated in *Ae. aegypti* (4). Ornamentation of midfemur (II): pale scales restricted to basal area in *Ae. albopictus* (5), entirely covered with pale scales in *Ae. Aegypti* (6). Ornamentation of clypeus: without silver scales, entirely black in *Ae. albopictus* (7), silvery white scales in one or two patches in *Ae. aegypti* (8).

**Figure 7.**
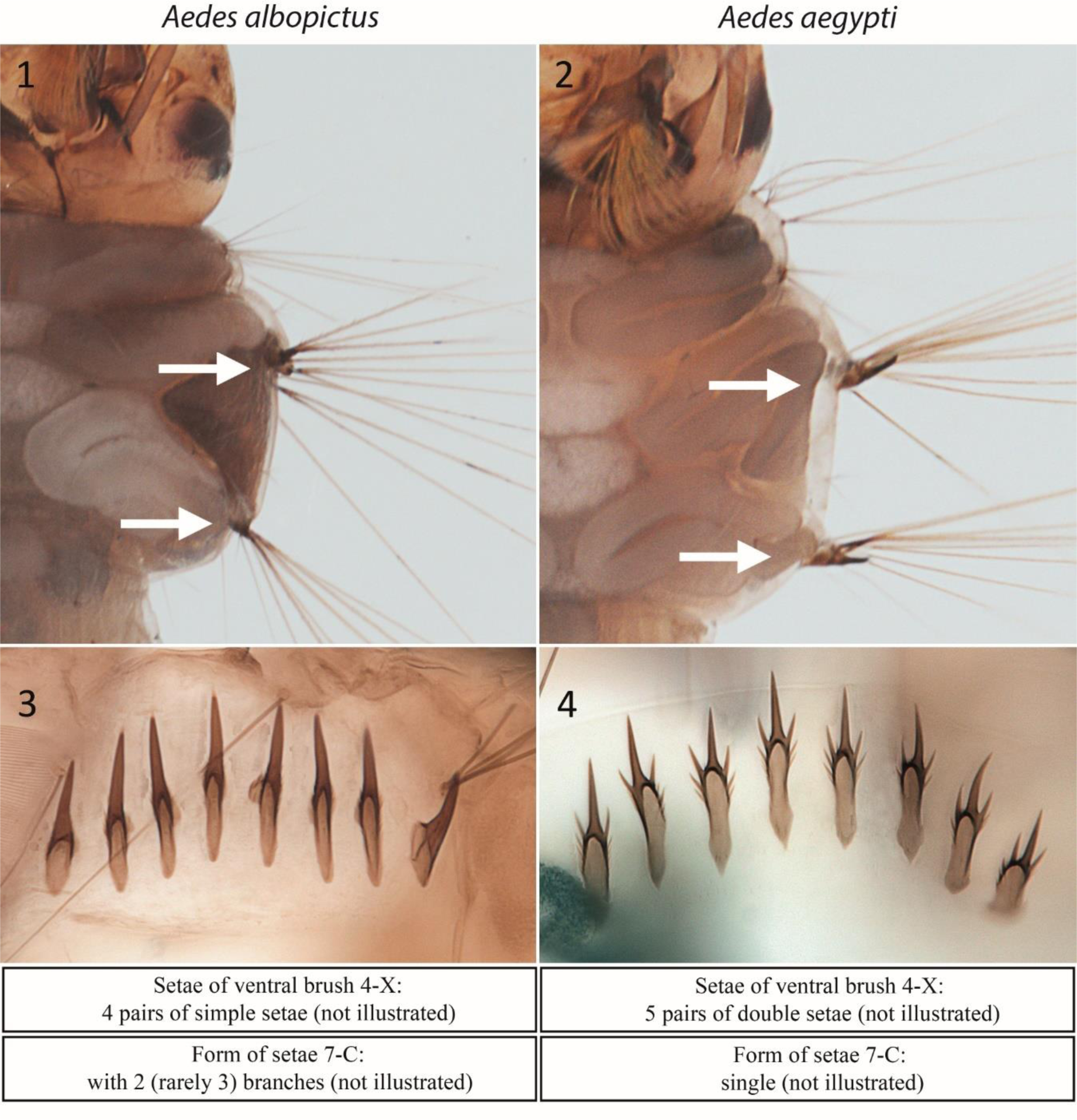
Morphological characteristic used to distinguish between *Ae. albopictus* and *Ae. aegypti* in L4 larvae. Specimens are from sample FLX-2023-021 of Rosario del Yata. Ornamentation of lateral support plates of setae 11-M and 11-T of meso- and metathorax: with tiny small spines in *Ae. albopictus* (1), and with large well-marked curved spines in *Ae. aegypti* (2). Form of the median comb scale of segment VIII: spine very long and sharp, with tiny lateral spines on base of scale in *Ae. albopictus* (3), and strong submedian spines (trident shape) in *Ae. aegypti* (4).

### Molecular identification

For *Ae. albopictus*, morphological identification was supplemented with molecular analysis using two mitochondrial gene fragments: Cytochrome C oxidase subunit 1 (COX1, also known as COI) and Cytochrome B (CYTB). Three larvae from Rosario del Yata (from samples FLX-2023-021, FLX-2023-043 and FLX-2023-044), and three from San Agustin (from samples FLX-2023-033B, FLX-2023-033C and FLX-2023-040) were used. DNA extraction followed a protocol previously proposed (^28^), based on CTAB/chloroform/isopropanol chemistry. Primers for both gene fragments were designed by one of us (CB). For COX1: Albo_Cox1_F: 5’- ACA AAT CAT AAA GAT ATT GGA ACA-3’ and Albo_Cox1_R : 5’- AAC TTC TGG ATG ACC AAA AA-3’; and for CYTB: Albo_CytB_F: 5’- TCA GCC TGA AAT TTT GGA-3’ and Albo_CytB_R : 5’- CAG GTT GAA TAT CAG GAG T -3’. PCR reactions were performed in 25µL final volume including 2µL of DNA (diluted at 20ng/µL), 1x buffer B (Hot FIREPol®), 1.5mM of MgCL2 (Hot FIREPol®), 0.8mM of dNTP (20mM) (Eurogentec), 5pmol of each primer and 1 Unit of FIREPol Taq DNA polymerase (Hot FIREPol®). PCR amplifications were carried out in Vapo Protect Thermocycler®. Cycling conditions were an initial denaturation at 95°C for 15min, followed by 35 cycles of 15s denaturation at 94°C, 30s annealing at 50°C for COX1 primers and 30s annealing at 53°C for CYTB primers and 1min extension at 72°C for both primers, and a final extension step at 72°C for 5min. The amplicons were sequenced by Azenta services, and the resulting sequences were manually corrected from the Sanger-type chromatograms. Subsequently, they were compared to deposited sequences in GenBank at NCBI using the basic local alignment search tool (BLAST) algorithm. Sequences returning a query coverage of >98% were selected to ensure removal of excessively short reference sequences. These selected sequences were then parsed using VSEARCH clustering commands (^29^) to identify all closely related haplotypes to our own sequences. Subsequently, maximum likelihood trees (ML trees) were constructed using the MEGA7 program (^30^) employing the best model of multiple substitutions suited to the data. The robustness of the clustering was assessed via a bootstrap procedure consisting of 100 replicates (^31^).

### Data preservation and storage

In the field, essential ecological data including GPS coordinates, types of larval breeding sites, and water quality were collected and organized using a cellular phone equipped with the VECTOBOL database (https://vectobol.ird.fr). The database was updated whenever an internet connection was available. The database was designed with REDCap electronic data capture tools hosted at IRD-France ^(32; 33)^. Then, in the laboratory, once species identifications were completed for each sample, the database was updated.

All collected samples have been deposited in the collections of the Medical Entomology Laboratory (LEMUMSS) at the Universidad Mayor de San Simón in Cochabamba, Bolivia. Some vouchers of *Ae. albopictus* were also deposited in the Entomology Laboratory of the Instituto Nacional de Laboratorios de Salud (INLASA) of the Ministry of Health in La Paz, Bolivia. In both collections, the sample’s IDs (which also correspond to the specimen prefix in collection) align with those listed in Table 1.

The DNA sequences of *Ae. albopictus* were deposited in GenBank with numbers PP465543 to PP465547 for COX1 and PP471874 to PP471878 for CYTB.

## RESULTS

### Collected species

In the peridomestic environment, *Ae. albopictus* was present in six of the 14 sampled houses, three in Rosario del Yata and three in San Agustin. The survey resulted in the identification of a total of 19 positive larval breeding sites among which six were positive for *Ae. albopictus*. In the field, the species was easily distinguishable by its unique morphological traits once it reached the adult stage (Fig. 6 and Fig. 8). During the survey, one *Ae. albopictus* female was captured while biting one of our team members (RTL) during larvae collection at sample ID: FLX-2023-044B (Table 1). Apart from *Ae. albopictus*, eight other mosquito species were collected from larval breeding sites in the peridomestic environment. They were: *Ae. aegypti* (seven instances)*, Ae. fluviatilis* (seven instances)*, Culex corniger* (one instance)*, Cx. coronator* (six instances)*, Cx. declarator* (four instances)*, Cx. quinquefasciatus* (six instances)*, Psorophora albigenu* (two instances) and *Toxorhynchites haemorrhoidalis* (three instances). As far as species associations are concerned, *Ae. albopictus* was collected in isolation in two instances, while in four instances, it was found in association with *Ae. aegypti* either alone (one instance), or also with *Cx. declarator* (one instance), or *Tx. haemorrhoidalis* (two instances) (Table 1). *Aedes aegypti* was found alone in three instances. The remaining ten collections with no *Stegomyia* consisted in *Cx. coronator, Cx. corniger, Cx. declarator, Cx. quinquefasciatus, Ae. fluviatilis* and *Ps. albigenu* in isolation or association (Table 1).

**Figure 8.**
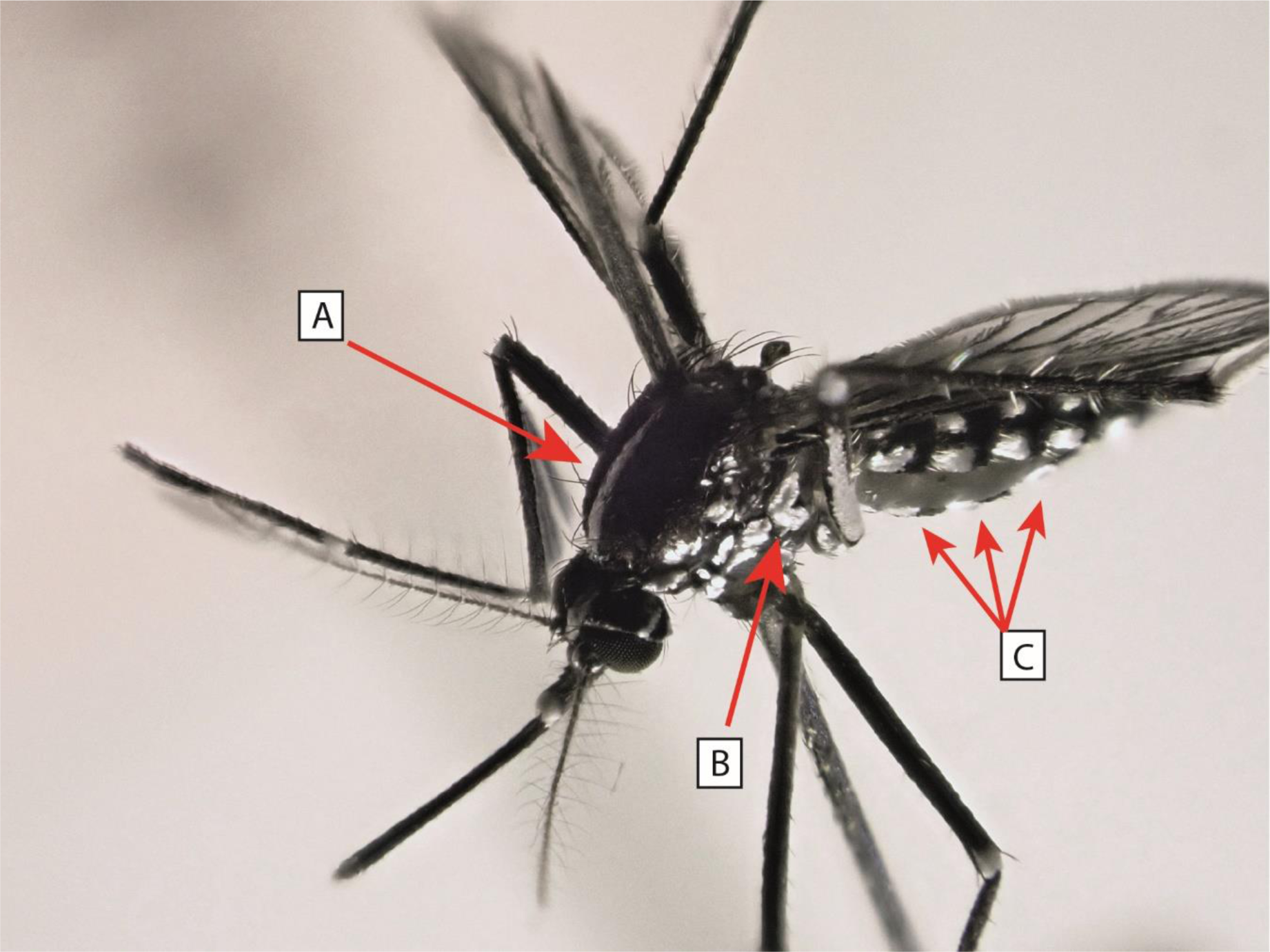
*Aedes albopictus* female (field collection ID: FLX-2023-021), reared from larvae, displaying distinctive species characteristics. These include a narrow white median longitudinal stripe against a black background on the scutum (A), two white scale spots forming a unique V shape on the mesepimeron (B), and the presence of pale scales exclusively at the basal part of each sternite III-V, resulting in an alternating pattern of white and black bands on the ventral surface (C).

In the “close to forest” environment and in the riverbank samples, no *Ae. albopictus* was collected. These two environments share with the peridomestic environment *Cx. coronator, Ae. fluviatilis* and *Ps. albigenu*. In the “close to forest” environment of Rosario del Yata, as in the peridomestic environment, *Cx. declarator* and *Tx. haemorrhoidalis* were also collected. In the “river environment”, various other species were also collected, including *Cx. pilosus, Cx. glyptosalpinx*, one unidentified *Culex sp*. and one *Anopheles sp*. (the two larvae too small and deteriorated to be identified) (Table 1).

Human bait collections in the peridomestic environment (by RTL), included one *Ae. albopictus* and one *Mansonia titillans* captured at 16:00 in Rosario del Yata. Meanwhile, in the ‘close to forest’ environment, captures carried out during one hour at the beginning of the night (FL, PB and RTL) consisted of numerous specimens of *Ma. titillans*. (Table 1).

### Molecular identification

The morphological identifications of *Ae. albopictus* were strongly supported by the molecular results. Specifically, for CYTB, the sequences obtained from five isolates precisely matched 46 reference sequences in GenBank, including accession number MN513368.1, as depicted in the ML tree (Fig. 9). Notably, this particular CYTB haplotype was detected across various countries such as Albania, Brazil, China, France, Greece, Italy, Japan, Portugal, and the United States. The tree analysis revealed *Ae. flavopictus* as the most closely related sister clade to *Ae. albopictus*.

**Figure 9.**
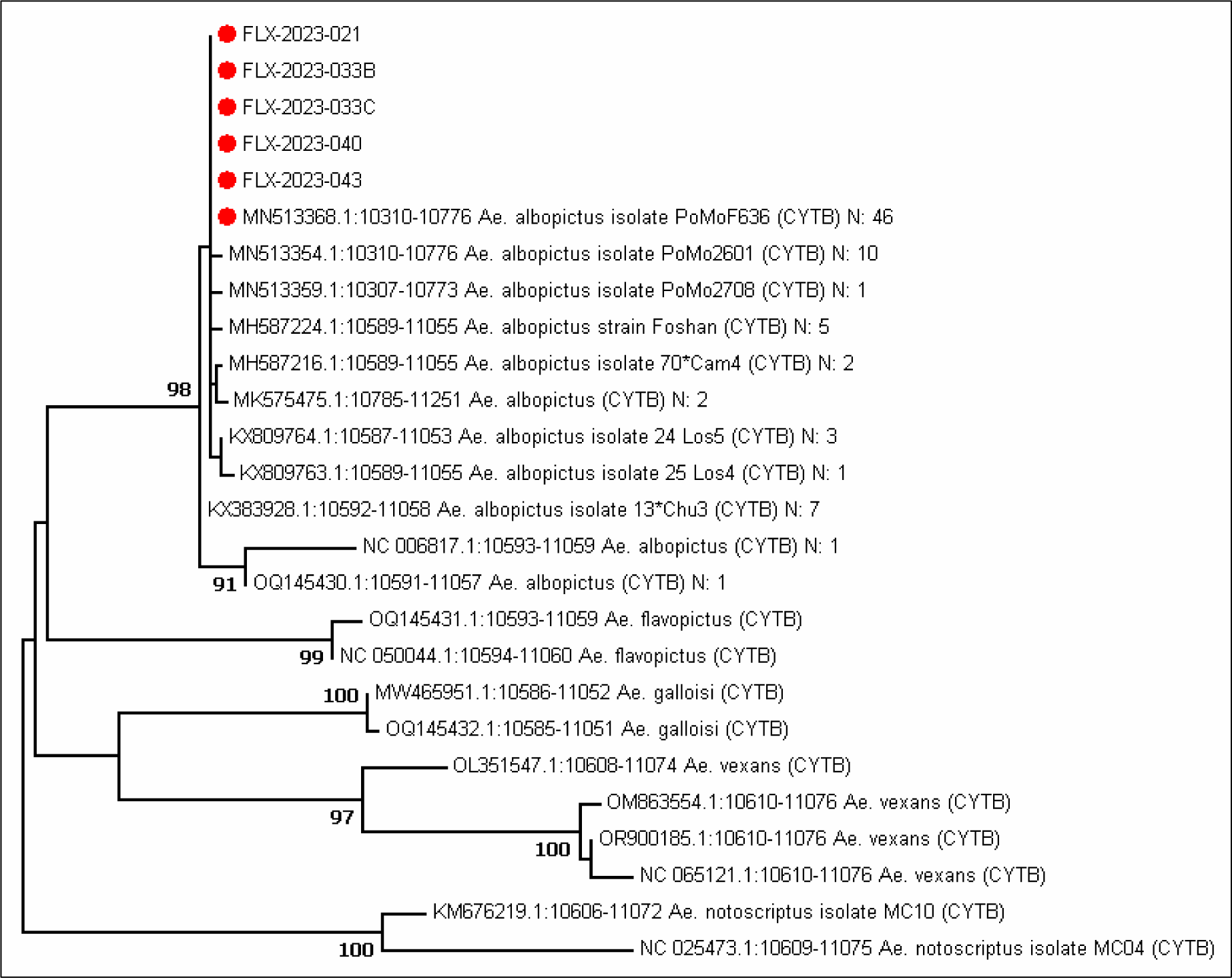
Maximum likelihood (ML) tree of 26 different CYTB haplotypes constructed from an alignment of 467 nucleotide positions. The numbers at the nodes correspond to the bootstrap values > 70%. The best multiple substitution model was the Tamura-Nei model +G (Gamma distribution). Our five isolates exactly matching with the GenBank reference MN513368.1 are visualized as red circles. N is the abundance of the particular *Ae. albopictus* haplotype in GenBank at that time.

Regarding COX1 (also known as COI), the sequences obtained from five isolates also exhibited an exact match with 564 reference sequences in GenBank, including accession number HQ398900.1, visualized in the ML tree (Fig. 10). In line with CYTB, this specific COX1 haplotype was recorded in multiple countries, including Brazil, where infested regions border Bolivia. This new ML tree further confirmed *Ae. flavopictus* as the most closely related species to *Ae. albopictus*.

**Figure 10.**
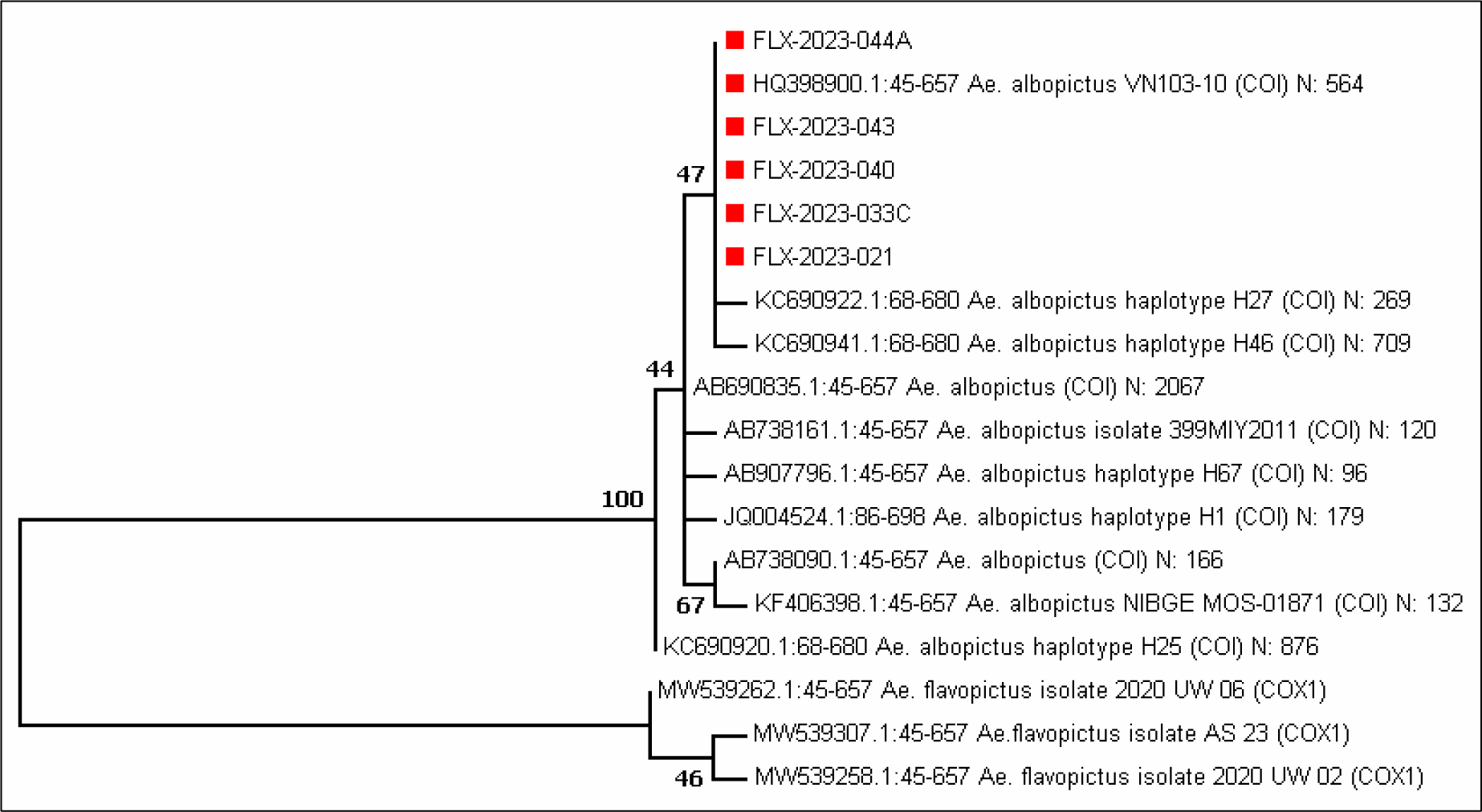
Maximum likelihood (ML) tree of 18 different COX1 (also called *COI*) haplotypes constructed from an alignment of 613 nucleotide positions. The numbers at the nodes correspond to the bootstrap values > 70%. The best multiple substitution model was the Tamura 3-parameter model +G (Gamma distribution). Our five isolates exactly matching with the GenBank reference HQ398900.1 are visualized as red squares. N is the abundance of the particular *Ae. albopictus* haplotype in GenBank at that time.

## DISCUSSION

The current study definitively identifies *Ae. albopictus* in diverse locations within Rosario del Yata and San Agustin, employing both standard morphological characteristics and molecular biology (COXI and CytB sequencing). In the field, more than 300 *Ae. Albopictus* larvae were collected of which more than 100 were reared until the adult stage. Moreover, one adult female was captured by chance in the field. This discovery, duly documented, puts an end to anecdotes that reported the species without substantiating evidence, a factor that has persistently generated doubt in numerous scientific articles.

The source of infestation in Rosario del Yata and San Agustin remains elusive. The presence of widespread haplotypes across both gene fragments precluded the identification of a potential origin for the initial insects colonizing the area. While sequencing additional genes might provide insight, the identical haplotypes observed in Brazil for both COX1 and CYTB suggest a plausible Brazilian origin for these insects as the most parsimonious hypothesis. It’s noteworthy that molecular identification effectively distinguishes *Ae. albopictus* from *Ae. flavopictus*, a closely related species that can be morphologically confused with *Ae. albopictus* (^34^).

The time of arrival is also unknown; however, it may be close to the current discovery date. Indeed, for years, various studies on *Ae. aegypti* in Bolivia have consistently failed to mention the collection of *Ae. albopictus*, despite the potential cohabitation of these two species in larval breeding sites, and the expectation of finding the latter species. This is particularly noteworthy given the extensive larval sampling efforts since 2005 at the country level, resulting in the collection of tens of thousands of samples ^(35; 36; 37; 38; 39; 40; 41)^. Furthermore, although the GBIF database does not display any collections of *Ae. albopictus*, it readily presents collection points for *Ae. aegypti* (^42^). Two of the article authors (FL and PB) did not find *Ae. albopictus* in 2022 during a similar entomological survey carried out in southern Beni, although they easily collected *Ae. aegypti* in eight occasions in a total of 15 peridomestic larval sites sampled in the Municipios of Palos Blancos, San Borja, Trinidad, Ascensión de Guarayos and Okinawa Uno. During the present 2023 survey, excluding the localities of Rosario del Yata and San Agustin, *Ae. albopictus* was not found, while *Ae. aegypti* was readily captured in seven out of the 12 peridomestic larval breeding sites sampled in the cities of Guayaramerin, Cachuela Esperanza and Inicua, suggesting a higher occurrence of the latter and, consequently, a more recent establishment of *Ae. albopictus*. The sampled localities mentioned in the previous references and the GBIF dataset(^42^) are depicted in Figure 11, illustrating the extensive geographical distribution of *Ae. aegypti* collecting points across Bolivia and its frequent occurrence.

**Figure 11.**
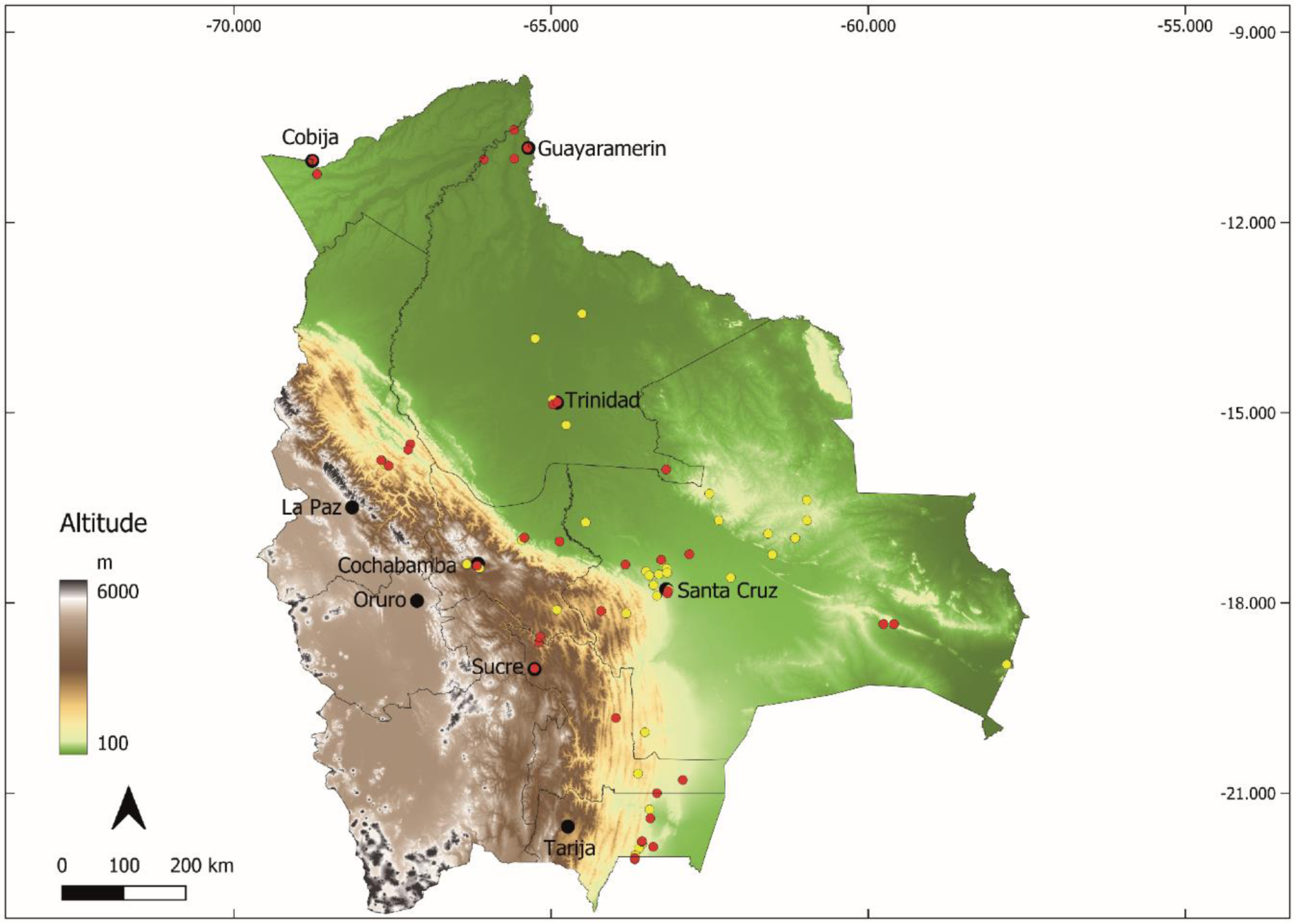
Occurrences of *Ae. aegypti* in Bolivia, represented by recent bibliographic references cited in the text (red dots) and the GBIF dataset (yellow dots). The distribution highlights the species’ frequent collection, contrasting with the absence of references for *Ae. albopictus* until the present study.

In the event of introduction via the tire or plant trade, the date of arrival in the country is likely to be close to the date of actual discovery, as predicted by a model suggesting, for Bolivia, a timeframe of 3 to 5 years from 2023 (^43^).

*Aedes albopictus* displays phenotypic plasticity, enabling the development of strains adapted to a broad range of environmental conditions and optimal temperatures for survival and development ^(44; 45)^. The mosquito has been collected along altitudinal transects, reaching up to 2100m above sea level in Nepal (^46^), and is commonly found above 1200 m in tropical environments ^(47; 48)^. With a demonstrated capacity to outcompete *Ae. aegypti* in diverse ecological contexts (^49^) and given the detection of *Ae. aegypti* in Bolivia’s major cities, including cities above 2000 m such as Sucre at 2700 m (^50^) and Cochabamba at 2600 m (^51^), it is probable that *Ae. albopictus* could eventually establish itself in these urban areas.

## CONCLUSION

When established in a new habitat, *Ae. albopictus* becomes of significant epidemiological concern, serving as a confirmed vector for the chikungunya virus and dengue virus, among others. The multifaceted topography and climate of Bolivia give rise to discrete ecosystems that intricately shape the prevalence and distribution of diverse insect vectors. Given the likelihood of *Ae. albopictus* colonizing regions where *Ae. aegypti* is already present, covering approximately 2/3 of the territory below 2700 meters in altitude, it becomes imperative for the Health Services overseeing vector surveillance to swiftly develop a situational map of *Ae. albopictus* occurrence in the country. This includes monitoring the species colonization process and incorporating *Ae. albopictus* into the targeted species for control measures. This proactive approach is crucial for effective public health management and the prevention of potential disease outbreaks associated with this invasive mosquito species.

## ACKNOWLEDGEMENTS

We express our gratitude to the residents of Rosario del Yata and San Agustin for granting us access to their backyards for sampling peridomestic larval breeding sites.

## AUTHORS’ CONTRIBUTION

Designing research study: FL

Funding Acquisition: FL, PB, LGO

Conducting experiments and acquiring data: FL, PB, RTL, AB, CB

Analyzing data: FL, PB, AB, CB

Writing – Original Draft: FL, PB, CB, RTL, AB, LGO

Review & Editing: FL, PB, CB, RTL, AB, LGO

## REFERENCES

1. Reinert JF, Harbach RE, Kitching IJ 2004. Phylogeny and classification of Aedini (Diptera: Culicidae), based on morphological characters of all life stages. Zool. J. Linn. Soc., 142, 289–368.

2. Lowe S, Browne M, Boudjelas S, De Poorter M 2000. *100 of the world’s worst invasive alien species. A selection from the Global Invasive Species Database*. The Invasive Species Specialist Group (ISSG), Auckland, New Zealand.

3. Lounibos LP 2002. Invasions by insect vectors of human disease. Annu. Rev. Entomol., 47, 233–266.

4. Adhami J, Reiter P 1998. Introduction and establishment of *Aedes (Stegomyia) albopictus* Skuse (Diptera: Culicidae) in Albania. J. Am. Mosq. Control Assoc., 14, 340–343.

5. Kraemer MUG, Sinka ME, Duda KA, Mylne A, Shearer FM, Brady OJ, Messina JP, Barker CM, Moore CG, Carvalho RG, Coelho GE, Van Bortel W, Hendrickx G, Schaffner F, Wint GRW, Elyazar IRF, Teng HJ, Hay SI 2015a. The global compendium of *Aedes aegypti* and *Ae. albopictus* occurrence. Scientific Data, 2.

6. Forattini OP 1986. *Aedes (Stegomyia) albopictus* (Skuse) identification in Brazil. Rev. Saúde Públ., 20, 244–245.

7. Garcia-Rejon JE, Navarro JC, Cigarroa-Toledo N, Baak-Baak CM 2021. An updated review of the invasive *Aedes albopictus* in the Americas; Geographical distribution, host feeding patterns, arbovirus infection, and the potential for vertical transmission of Dengue virus. Insects, 12.

8. Torales Ruotti M 2017. *Plan de manejo integrado de vectores.* Ministerio de Salud Publica y Bienestar Social, Paraguay, 64 pp.

9. Goenaga S, Chuchuy A, Micieli MV, Natalini B, Kuruc J, Kowalewski M 2020. Expansion of the distribution of *Aedes albopictus* (Diptera: Culicidae): New records in Northern Argentina and their implications from an epidemiological perspective. J. Med. Entomol., 57, 1310–1313.

10. Pancetti FGM, Honório NA, Urbinatti PR, Lima-Camara TN 2015. Twenty-eight years of *Aedes albopictus* in Brazil: a rationale to maintain active entomological and epidemiological surveillance. Rev. Soc. Bras. Med. Trop., 48, 87–89.

11. Costa Rocha R, Silva Cardoso A, Lunier de Souza J, Silva Pereira E, Fernandes de Amorim M, Martins de Souza MS, Lima Medeiros C, Mendes Monteiro MF, Oliveira Meneguetti DU, Bicudo de Paula M, Brilhante AF, Nunes Lima-Camara T 2023. First official record of *Aedes (Stegomyia) albopictus* (Diptera: Culicidae) in the Acre State, Northern Brazil. Rev. Inst. Med. Trop. São Paulo, 65, e20.

12. Reiter P 1998. *Aedes albopictus* and the world trade in used tires, 1988-1995: The shape of things to come? J. Am. Mosq. Control Assoc., 14, 83–94.

13. Benedict MQ, Levine RS, Hawley WA, Lounibos LP 2007. Spread of the tiger: Global risk of invasion by the mosquito *Aedes albopictus*. Vector Borne Zoonotic Dis., 7, 76–85.

14. Pan American Health Organization 1996. Study on the feasibillity of erradication Aedes aegypti, CE118/16, 1–17.

15. Pan American Health Organization 1997. The feasibility of eradicating *Aedes aegypti* in the Americas. Pan American Journal of Public Health, 1, 68–72.

16. Serrano TI 2023. Descubren nuevo y más agresivo mosquito transmisor de: dengue, zika, chikunguña y fiebre amarilla en Santa Cruz. El Deber.

17. Ibaraki F, Gómez L, Arteaga S 2023. Primer Registro de *Aedes albopictus* vector del Dengue, Zika y Chikungunya. Resultado de la investigación en el municipio de San Ignacio de Velasco. In, Facebook Video.

18. GBIF Occurrence download [database on the Internet]2024; unique ID: 10.15468/dl.v2c3fx.

19. Kraemer MUG, Sinka ME, Duda KA, Mylne AQN, Shearer FM, Barker CM, Moore CG, Carvalho RG, Coelho GE, Van Bortel W, Hendrickx G, Schaffner F, Elyazar IRF, Teng HJ, Brady OJ, Messina JP, Pigott DM, Scott TW, Smith DL, Wint GRW, Golding N, Hay SI 2015b. The global distribution of the arbovirus vectors *Aedes aegypti* and *Ae. albopictus*. ELIFE, 4.

20. Harrison BA, Byrd BD, Sither CB, Whitt PB 2016. *The Mosquitoes of the Mid-Atlantic Region: An Identification Guide*. Western Carolina University, Madison Height, MI, 201 pp.

21. Rossi G, Martinez M 2013. Lista de especies y clave ilustrada para la identificación de larvas de mosquitos (Diptera : Culicidae) halladas criando en recipientes artificiales en Uruguay. Bol. Soc. Zool. Urug., 22, 49–65.

22. Darsie RF, Ward RA 2005. *Identification and geographical distribution of the mosquitos of North America, North of Mexico*. University Press of Florida, 384 pp.

23. Huang YM 1968. Neotype designation for *Aedes (Stegomyia) albopictus* (Skuse) (Diptera: Culicidae). Proc. Entomol. Soc. Wash., 70, 297–302.

24. Huang YM 1979. Medical entomology studies – XI. The subgenus *Stegomyia* of *Aedes* in the Oriental region with keys to the species (Diptera: Culicidae). Contrib. Am. Entomol. Inst., 15, 1–79.

25. Savage HM, Smith GC 1994. Identification of damaged adult female specimens of *Aedes albopictus* and *Aedes aegypti* in the New World. J. Am. Mosq. Control Assoc., 10, 440–442.

26. Tanaka K, Mizusawa K, Saugstad ES 1979. A revision of the adult and larval mosquitoes of Japan (including the Ryukyu Archipelago and Ogasawara Islands) and Korea (Diptera: Culicidae). Contrib. Am. Entomol. Inst., 16, 1–987.

27. Darsie RJJ 1985. Mosquito of Argentina. Part I. Keys for identification of adults females and fourth stage larvae in English and Spanish (Diptera, Culicidae). Mosquito Systematics, 17, 153–253.

28 . Berger A, Le Goff G, Bousses P, Rahola N, Ferre JB, Ayala D, Robert V 2022. Using a pupal exuvia to designate the undamaged neotype of a species belonging to a complex of sibling species - the case of *Aedes coluzzii* (Diptera, Culicidae). Parasite, 29, 19.

29. Rognes T, Flouri T, Nichols B, Quince C, Mahe F 2016. VSEARCH: a versatile open source tool for metagenomics. PeerJ, 4, e2584.

30. Kumar S, Stecher G, Tamura K 2016. MEGA7: Molecular Evolutionary Genetics Analysis Version 7.0 for Bigger Datasets. Mol. Biol. Evol., 33, 1870–1874.

31. Felsenstein J 1985. Confidence-Limits on Phylogenies - an Approach Using the Bootstrap. Evolution, 39, 783–791.

32. Harris PA, Taylor R, Thielke R, Payne J, Gonzalez N, Conde JG 2009. Research electronic data capture (REDCap)--a metadata-driven methodology and workflow process for providing translational research informatics support. J Biomed Inform, 42, 377–381.

33. Harris PA, Taylor R, Minor BL, Elliott V, Fernandez M, O’Neal L, McLeod L, Delacqua G, Delacqua F, Kirby J, Duda SN, Consortium RE 2019. The REDCap consortium: Building an international community of software platform partners. J Biomed Inform, 95, 103208.

34. Chaves LF, Friberg MD 2021. *Aedes albopictus* and *Aedes flavopictus* (Diptera: Culicidae) pre-imaginal abundance patterns are associated with different environmental factors along an altitudinal gradient. Curr Res Insect Sci, 1, 100001.

35. Chucatiny CH 2013. Caracterización de haplotipos del gen ND4 en poblaciones de Aedes aegypti (vector del dengue) de las comunidades de San Borja y Caranavi. Licenciatura, Universidad Mayor de San Andrès, La Paz, 91 pp.

36. Salas R 2007. *Susceptibilidad y resistencia de Aedes aegypti a los insecticidas y mecanismos de resistencia en tres cepas de Bolivia 2006-*2007. Licenciatura, Universidad Autonoma Gabriel Moreno, Santa Cruz de la Sierra, Bolivia, 100 pp.

37. Valdéz Zamorano NG 2009. *Caracterización morfológica y genética de poblaciones urbanas y rurales de Aedes (Stegomyia) aegypti L. (Diptera - Culicidae) ubicadas en localidades endémicas de dengue en Bolivia*. Licenciatura, UAGRM, IRD, Santa Cruz de la Sierra (BOL), La Paz, 100 multigr. pp.

38. Lopez Rodriguez RW 2015. *Estudio de la sensibilidad y/o resistencia a los insecticidas del Aedes aegypti, vector del dengue en Bolivia*. Magister, Universidad Mayor de San Andrès, La Paz, Bolivia, 56 pp.

39. Paupy C, Le Goff G, Brengues C, Guerra M, Revollo J, Simon ZB, Herve JP, Fontenille D 2012. Genetic structure and phylogeography of *Aedes aegypti*, the dengue and yellow-fever mosquito vector in Bolivia. Infection Genetics and Evolution, 12, 1260–1269.

40. Barja-Simon Z, Le Goff G, Callata R, Walter A, Bremond P 2009. Infestación de los cementerios de Santa Cruz de la Sierra por los mosquitos vectores del dengue. Rev. Enferm. Infecc.Trop., 1, 29–32.

41. Bremond P, Roca Y, Breniere SF, Walter A, Barja-Simon Z, Fernandez RT, Vargas J 2015. Evolution of dengue disease and entomological monitoring in Santa Cruz, Bolivia 2002 - 2008. PLoS One, 10, e0118337.

42. GBIF. Org User 2024. Occurrence Download. In, The Global Biodiversity Information Facility.

43. Oliveira S, Capinha C, Rocha J 2023. Predicting the time of arrival of the Tiger mosquito (*Aedes albopictus*) to new countries based on trade patterns of tyres and plants. J. Appl. Ecol., 60, 2362–2374.

44. Waldock J, Chandra NL, Lelieveld J, Proestos Y, Michael E, Christophides G, Parham PE 2013. The role of environmental variables on *Aedes albopictus* biology and chikungunya epidemiology. Pathogens and Global Health, 107, 224–241.

45. Kramer IM, Pfeiffer M, Steffens O, Schneider F, Gerger V, Phuyal P, Braun M, Magdeburg A, Ahrens B, Groneberg DA, Kuch U, Dhimal M, Müller R 2021. The ecophysiological plasticity of *Aedes aegypti* and *Aedes albopictus* concerning overwintering in cooler ecoregions is driven by local climate and acclimation capacity. Sci. Total Environ., 778.

46. Dhimal M, Gautam I, Joshi HD, O’Hara RB, Ahrens B, Kuch U 2015. Risk factors for the presence of Chikungunya and Dengue vectors (*Aedes aegypti* and *Aedes albopictus)*, their altitudinal distribution and climatic determinants of their abundance in Central Nepal. PLOS Neglected Tropical Diseases, 9.

47. Delatte H, Dehecq JS, Thiria J, Domerg C, Paupy C, Fontenille D 2008. Geographic distribution and developmental sites of *Aedes albopictus* (Diptera: Culicidae) during a Chikungunya epidemic event. Vector Borne Zoonotic Dis., 8, 25–34.

48. Sayono S, Nurullita U, Sumanto D, Handoyo W 2017. Altitudinal distribution of *Aedes* indices during dry season in the dengue endemic area of Central Java, Indonesia. Annals of Parasitology, 63, 213–221.

49. Lounibos LP, Juliano SA 2018. Where Vectors Collide: The Importance of Mechanisms Shaping the Realized Niche for Modeling Ranges of Invasive *Aedes* Mosquitoes. Biol. Invasions, 20, 1913–1929.

50. Ríos C, Rosas N, Delgadillo-Iglesias MA, Solis-Soto MT 2023. Presence of *Aedes aegypti* in a high-altitude area in Bolivia. bioRxiv, 2023.2008.2007.552199.

51. Aquino Rojas E, Rojas Cortez M, Espinoza J, Vallejo E, Lozano D, Torrico F 2016. Caracterización de la infestación de viviendas por *Aedes aegypti* en el área metropolitana de Cochabamba, Bolivia: nuevos registros altitudinales. Gac. Med. Bol., 39, 83–87.

